# Addressable and adaptable intercellular communication via DNA messaging

**DOI:** 10.1101/2022.11.17.516988

**Authors:** John P. Marken, Richard M. Murray

## Abstract

Engineered consortia are a major research focus for synthetic biologists because they can implement sophisticated behaviors inaccessible to single-strain systems. However, this functional capacity is constrained by their constituent strains’ ability to engage in complex communication. DNA messaging, by enabling information-rich channel-decoupled communication, is a promising candidate architecture for implementing complex communication. But its major advantage, its messages’ dynamic mutability, is still unexplored. We develop a framework for addressable and adaptable DNA messaging that leverages all three of these advantages and implement it in a plasmid conjugation-based communication channel. Our system can bias the transfer of messages to targeted receiver strains by 100-to 1000-fold, and their recipient lists can be dynamically updated *in situ* to control the flow of information through the population. This work lays the foundation for future developments that further utilize the unique advantages of DNA messaging to engineer previously-inaccessible levels of complexity into biological systems.

## Introduction

A major current focus of synthetic biology research is to expand beyond the the field’s original paradigm of engineering a single cell strain for a particular application and to instead engineer consortia, which are populations consisting of multiple distinct cell types [1, 2]. By enabling the division of labor among its constituent strains, a consortia-based approach allows each strain to specialize itself to its assigned task while minimizing the metabolic burden to itself [3]. Engineered consortia are therefore able to achieve higher levels of functional complexity [4, 5, 6] and evolutionary stability [7, 8] than analogous single-strain systems.

In order for an engineered consortium to function properly, however, it is necessary that each of its constituent strains can stably coexist and act in concert with each other. This coordinated activity is maintained by intercellular communication systems that allow the strains to dynamically instruct each other to perform programmed functions, like modulating their growth rate or activating a target gene. The achievable complexity of a consortium’s behavior is therefore constrained by the capacity of its communication channels to transmit complex messages [6]. Realizing this, the synthetic biology community has placed much effort towards expanding the toolbox of intercellular communication channels and enabling increasingly information-dense communication between cells [9, 10, 11, 12, 13, 14, 15].

These efforts have almost exclusively focused on a molecular architecture that we will term Small Molecule Actuated communication (SMA communication), wherein a sender cell synthesizes a small molecule that diffuses through the extracellular environment to enter a receiver cell that contains the requisite machinery to initiate a preprogrammed response to the signal. SMA communication channels were originally implemented using molecular parts co-opted from quorum sensing systems [9], but in recent years the toolbox has expanded to include metabolites [10], hormones [13], and antibiotics [7] as signal vectors.

An alternative molecular architecture, DNA messaging, was proposed in a pioneering report by Ortiz and Endy [14]. Here, horizontal gene transfer mechanisms are co-opted into a communication channel that transmits DNA-encoded messages between cells (Fig S1). Because the actual content of the message is an arbitrary genetic sequence within the mobile vector itself, Ortiz and Endy coined the term “message-channel decoupling” to describe the fact that a single DNA-based communication channel can send different messages that contain different types of instructions to the recipient cells [14].

In contrast, SMA communication channels exhibit message-channel coupling because the nature of the encodable message is tied to the molecular identity of the signaling molecule— a homoserine lactone, for example, can only be used to encode the message “activate the cognate transcription factor”, and an antibiotic can only be used to encode the message “kill the susceptible cell strains” (Fig 1a).

**Figure 1.**
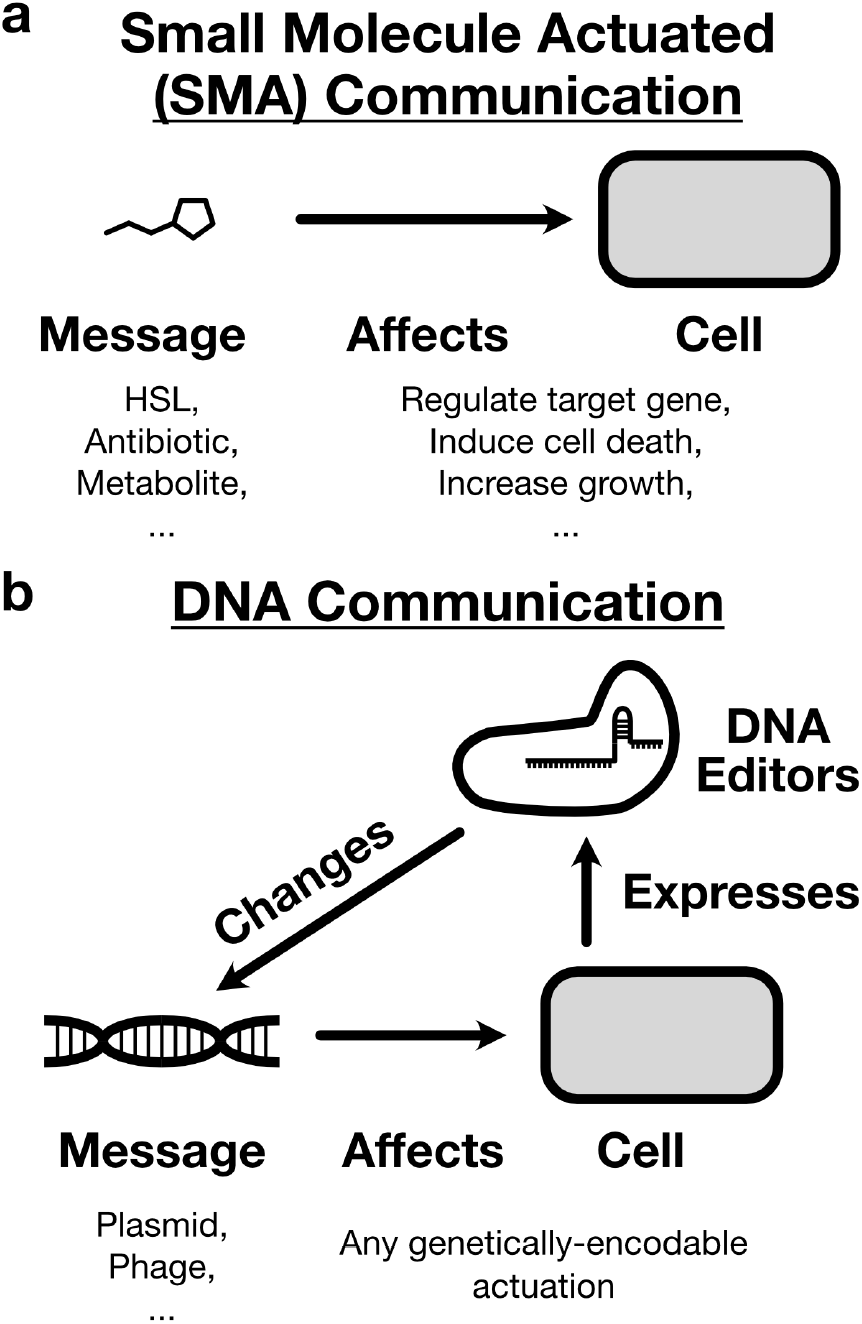
Schematics of different architectures for engineered intercellular communication. (a) Small Molecule Actuated (SMA) communication systems exhibit message-channel coupling, meaning that the behavior they induce in the receiver cell is hardcoded into the molecular identity of the signaling molecule itself. This molecular identity cannot be changed without disrupting the functioning of the channel itself. (b) DNA communication systems exhibit message-channel decoupling, meaning that a given channel can transmit multiple types of messages to induce any genetically-encodable response in the receiver cell. Furthermore, the cells themselves can express molecular DNA editors to change the content of the messages *in situ*, closing the loop to enable autonomous system reconfiguration.

A second important advantage of DNA communication is that a single DNA message can encode a large amount of information content, as many horizontal gene transfer systems can easily transfer several kilobases of arbitrary sequence [16, 17, 18]. In contrast, SMA channels can only modulate their activity via the concentration of their signal vector, a single small molecule. This heavily constrains the information density of the communication channel, to the point where in applications like digital computation where concentrations are interpreted binarily as either OFF or ON, a single SMA channel can only transmit a single bit of information [19].

Together, these advantages suggest that DNA messaging is an ideal communication architecture for engineering complex consortia with sophisticated information processing requirements. But in the ten years since its introduction, DNA messaging has not been widely adopted by synthetic biologists. In fact, although there have been some computational studies of various implementations [20, 21, 22] and horizontal gene transfer systems have been co-opted by synthetic biologists for engineering environmental microbiomes [23, 24, 25], to our knowledge the original Ortiz-Endy implementation remains the only reported experimental usage of DNA-based communication to date.

Why is the case? One reason is that, though it was pioneering in its foresight, the Oritz-Endy implementation did not demonstrate a third property of DNA communication that is critical in enabling the implementation of qualitatively new functionalities— the dynamic mutability of DNA messages. Unlike SMA channels, where the message is encoded into the structure of an immutable signal molecule, cells have the ability to express DNA editors that can make targeted changes to the content of the message *in situ* (Fig 1b). This ability has only expanded with the recent explosion in research on programmable DNA editors like CRISPR-Cas systems, integrases, and base editors [26, 27]. Although theoretical reports have rightly identified mutability as a key advantage of DNA messaging [20], to date this property has not been experimentally demonstrated.

We therefore set out to develop a general and scalable architecture for DNA messaging that allows users to fully take advantage of all three of its unique properties: message-channel decoupling, high information density, and dynamic message mutability. In order to ensure our framework’s compatibility with arbitrary messages transferred along arbitrary horizontal gene transfer systems, we used channel-orthogonal molecular tools to implement a functionality that is required in all communication systems— the ability to address the message to a targeted set of recipients.

Our addressing framework uses CRISPR-Cas systems to internally validate each message transfer event within the consortium, enabling the targeted delivery of a given message to any subset of the strains in a population. We additionally design a framework for using integrases to modularly update messages’ recipient lists *in situ*, enabling the control of information flow through a population. This work establishes a universally-applicable framework for effective DNA-based communication that sets the stage for future efforts that expand its ability to implement previously-inaccessible functionalities into engineered consortia.

## Results

### Incorporating message addressability into a plasmid conjugation-based communication system

We first describe the implementation of an addressability system for our DNA messaging framework. Any such implementation requires a means for the molecular recognition of specific genetic sequences, and we chose to use the CRISPR-Cas adaptive immunity system due to its ability to programmatically target and cleave desired nucleotide sequences on genetic vectors entering the cell [28, 29]. Although multiple different Cas systems have been demonstrated to cleave and degrade DNA vectors within cells [30, 31], we specifically chose to use the *S. pyogenes* Cas9 endonuclease system because it contains the required binding, unwinding, and cleaving activities within a single protein, facilitating its use in many different host organisms [32]. Additionally, well-developed procedures exist for generating large libraries of orthogonal single-guide RNAs (gRNAs) for the Cas9 system [15, 33], and the small footprint of the gRNA binding site (23 bp) means that many such sites can be incorporated onto a DNA message without significantly burdening any potential sequence length constraints from the transfer system. Together, these properties make the Cas9-gRNA system an ideal candidate for implementing a scalable, modular, and host-orthogonal addressing system for DNA messaging.

The design of our addressability framework is as follows. Each receiver cell in the consortium expresses both Cas9 and a unique gRNA that serves as a molecular signature encoding its strain identity. The sender cells themselves require no additional molecular machinery, but the DNA message must now contain an array of gRNA binding sites that correspond to the receiver strains that should *not* receive the message. This array is termed the “address region” because it acts as a blocklist, encoding the recipient list of the message as the set of strains whose gRNAs are *not* encoded in the address (Fig 2a). When the message is transferred to a receiver cell, the Cas9-gRNA complex checks the validity of the transfer— if the transfer is invalid, then the complex will bind to the cognate site on the address region and cleave the message, leading to its degradation. If the transfer is valid, then the complex does not interact with the message and it is able to freely propagate within the receiver cell. This process is schematized in Fig 2b.

**Figure 2.**
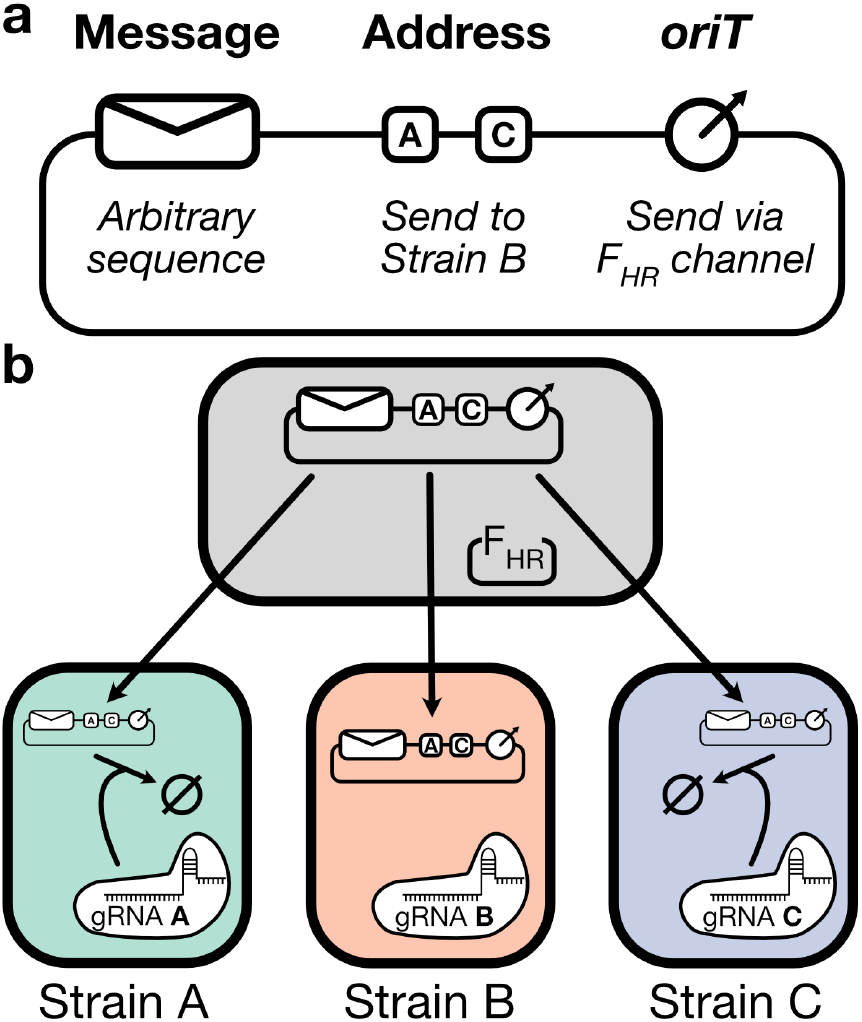
(a) Schematic of an addressable DNA message. The content of the message is an arbitrary genetic sequence, and the address region uses gRNA binding sites to act as a blocklist that determines the message’s recipient list by excluding transfer to all encoded strains. The origin of transfer (*oriT*) allows the message to interact with the cognate horizontal gene transfer machinery in the sender cell. (b) Schematic of transfer blocking. The DNA message is initially transferred promiscuously to all receiver strains in the population. As the message enters a receiver cell, the binding sites on the address region become exposed to cleavage by the Cas9-gRNA complex expressed within the cell. This cleavage only occurs if a binding site on the address matches the gRNA expressed in the receiver cell, thus ensuring that the message only persists within its appropriate recipients by eliminating the messages sent to invalid recipients.

An important property of this addressing framework is that the transfer validation system interacts with the message itself, rather than the transfer machinery that carries the message. This means that *a single DNA channel can send messages that are addressed to different recipients*. When addressability is implemented via channel-intrinsic properties, such as in the Ortiz-Endy system’s reliance on the M13 bacteriophage’s narrow infection host range [14], every message that is transmitted by a channel must go to the same recipient list regardless of its content.

In demonstrating the incorporation of our message addressing framework into a DNA-based communication system, we chose to deviate from Ortiz and Endy’s original choice of the filamentous bateriophage M13 and instead used a plasmid conjugation-based communication system. This is because the properties of plasmid conjugation systems are better-aligned with the advantages of DNA-based communication as a whole— plasmids can encode larger messages, with conjugative plasmids regularly reaching lengths of hundreds of kilobases [34, 18], and can transfer to taxonomically-diverse recipients [35, 36], facilitating their use in multispecies consortia. We specifically chose to use the F_HR_ system developed by Dimitriu et al. [16], which is based on the *E. coli* fertility factor F, the canonical representative of conjugative plasmids [37].

### Cas9-mediated blocking of plasmid receipt is inducible and orthogonal

In order to demonstrate that Cas9-mediated cleavage can indeed block the receipt of a mobilized plasmid, we performed pairwise sender-receiver experiments in the F_HR_ -based communication system. Receiver cells containing a genomically-integrated spectinomycin resistance cassette were transformed with a plasmid encoding OHC14-HSL-inducible expression of Cas9 and one of two gRNAs (“A” or “B”), and sender cells containing a genomically-integrated apramycin resistance cassette were transformed with the F_HR_ helper plasmid and a pSC101 message plasmid that constitutively expresses a chloramphenicol resistance gene. Two variants of this message plasmid were constructed, differing in whether their address region contained a single A binding site or a single B binding site (Fig 3a).

**Figure 3.**
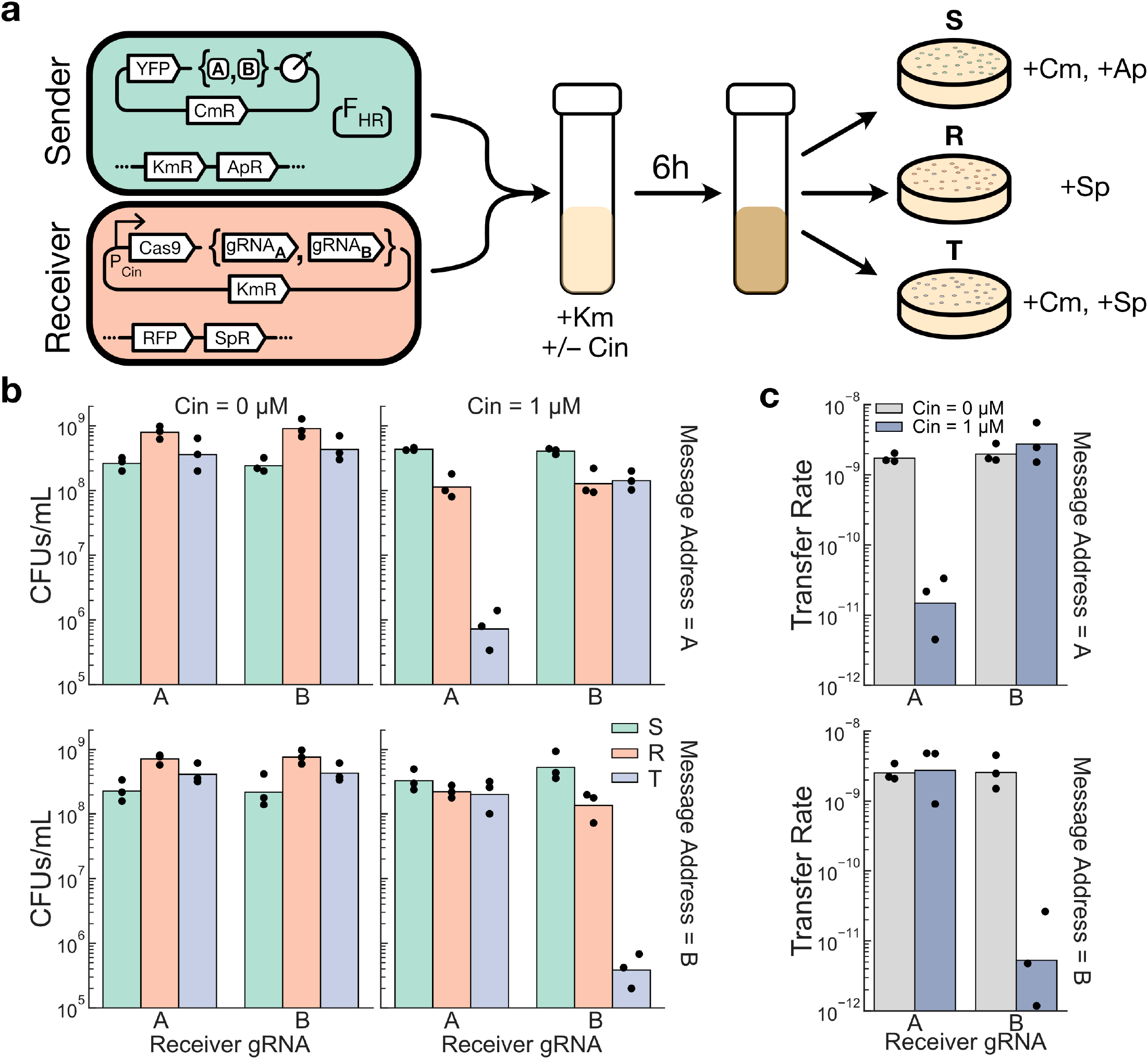
Cas9-mediated cleavage of incoming plasmids can bias their transfer to targeted recipients. (a) Schematic of the experimental setup. Senders (S) and Receivers (R) carrying one of two plasmid variants are grown together in a coculture and selective plating is used to isolate them, as well as the transconjugants (T), from the mixed culture. Note that transconjugants will appear on the receiver-selecting plates so R is the *total* density of receivers in the population (Methods). (b) Endpoint strain densities, measured in colony forming units (CFUs) per mL of culture. (c) Transfer rates, calculated as T/(S * R), of the message plasmid in each of the conditions in (b). Dots show the values from three biological replicates measured on different days, and bars depict the geometric mean of these values.

With this setup, selective plating could be used to individually isolate the senders, receivers, and transconjugants from a mixed population and calculate their densities. We performed mating experiments on all four combinations of sender-receiver pairs in the presence and absence of OHC14-HSL induction and measured the densities of each strain after 6 hours of growth in a shaken LB coculture (Fig 3b).

We then quantified the effectiveness of Cas9-mediated plasmid blocking by calculating the plasmid transfer rate in each experiment, defined as the transconjugant density divided by the product of the total sender and receiver densities. We observed that the A-containing message plasmid had a 185-fold higher transfer rate to its valid recipient (the B receiver) than to its invalid recipient (the A receiver), and that for the B-containing message plasmid the difference was 520-fold (Fig 3c). When the Cas9 system was not induced in the receiver cells, this biased transfer was not observed (Fig 3c).

Having demonstrated that our addressability system performed successfully in a two-strain population, we next asked whether our system could scale to multi-strain populations where a given address region may need to encode several gRNA binding sites. We constructed three different receiver strains that, in addition to the spectinomycin resistance gene, each express a distinct fluorescent protein (RFP, YFP, or BFP) from a genomically-integrated cassette. In this way, all three receivers could be mixed together with the sender strain in a four-strain coculture and the colors could be used to determine the density of each distinct receiver strain after selective plating. In order to further assess the generality of our Cas9-mediated blocking system, we used a set of orthogonal gRNAs developed by Didovyk et al. [33] instead of reusing the A and B gRNAs from the previous experiment. We transformed each of the colored receiver strains with a plasmid encoding Cas9 and one of three of the Didovyk gRNAs (D1, D2, or D3), and constructed sender strains containing one of eight message plasmids addressed to every possible combination of the three receiver strains (Fig 4a).

**Figure 4.**
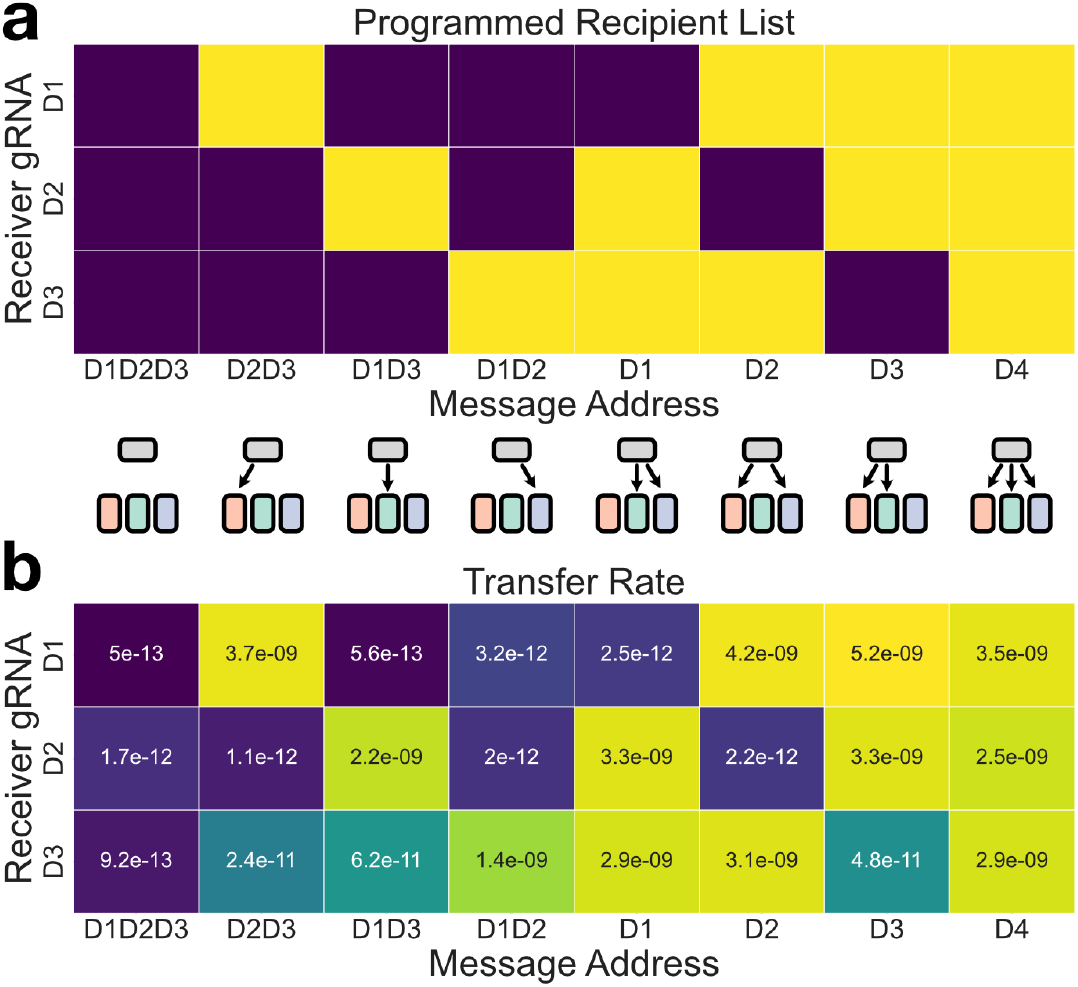
Programmable delivery of message plasmids to arbitrary subsets of a multi-strain population. (a) Schematic representation of the intended recipient list for each of the eight message plasmids. Dark squares indicate an invalid transfer and light squares indicate a valid transfer. (b) The observed geometric mean of the transfer rates to each receiver type, calculated from three biological replicates measured on different days. The color map is scaled logarithmically over four orders of magnitude. Individual transfer rate values are shown in Fig S2.

We found that even in the more complex setting of a four-strain population, our system was able to preferentially deliver the message to its appropriate recipients, often with a transfer rate that was over 1000-fold higher to valid recipients than to invalid recipients (Fig 4b; Fig S2). Although the three gRNAs used in the receivers were previously reported to be of comparable effectiveness in a dCas9-mediated transcriptional repression assay [33], the D1 and D2 gRNAs were able to block invalid transfers much more strongly than the D3 gRNA— the geometric mean of the fold change in transfer rates between valid and invalid recipients across all conditions where the invalid recipients expressed the D3 gRNA was 79-fold, compared to 1256-fold and 1577-fold for the D1 and D2 gRNAs, respectively (Fig S2).

### Cells can use integrases to edit DNA messages *in situ* and update their recipient list

Having demonstrated that our Cas9-mediated blocking system can successfully implement high-fidelity addressable communication between cells, we next proceeded to incorporate adaptability into the message transmission framework by enabling the programmable *in situ* editing of a message’s recipient list. This can be accomplished by applying molecular DNA editors to modify the gRNA binding sites on the address region.

Specifically, a system for programmable address editing should have the ability to both add a new binding site to the array and remove (or invalidate) an existing binding site from the array.

Serine integrases are a class of proteins that are well-suited for this task because of their ability to bind to specific attachment sequences and add, remove, or swap the regions between these sites depending on their configuration and orientation along the DNA [38]. Their efficiency and programmability have made their use ubiquitous among both molecular and synthetic biologists, and large sets of diverse and orthogonal integrases have been characterized [39, 40].

We implement address editing by flanking each binding site on the address region with orthogonal integrase attachment sites, in such a way that the expression of the cognate integrase will swap the binding site with a different binding site contained on a separate non-mobile plasmid via a process called recombinase-mediated cassette exchange [42] (Fig 5a). This procedure leaves the rest of the message, including the other binding sites on the address region, unaffected.

**Figure 5.**
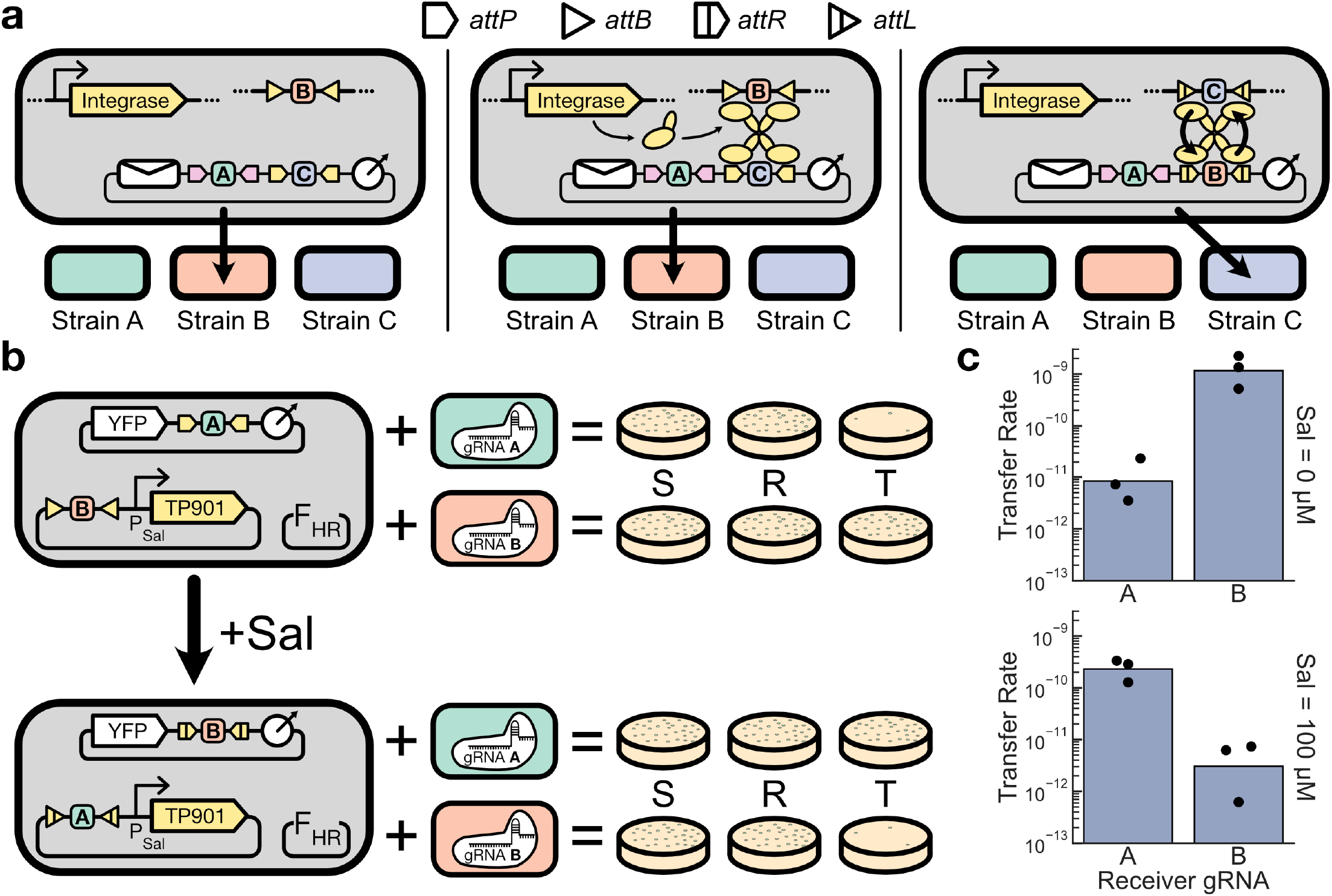
Integrase-mediated address editing. (a) Schematic of the process. In the left panel, expression of the integrase has not been induced and no editing has occurred. The message is addressed to Strain B. Orthogonal integrase attachment sites flank each binding site on the address region. In the middle panel, the integrase associated with the C site on the address region has been induced. The corresponding attachment sites for this integrase are also encoded separately on a sequence distinct from the message plasmid. In the right panel, the cassette exchange process has been completed, and the C site on the address region has been swapped with a B site, updating the message’s recipient list to Strain C. The process is unidirectional as it converts the attB and attP sites into attL and attR sites that can no longer undergo exchange, making this change permanent unless the cognate reverse directionality factors are expressed to reverse the process and restore the original sequence configuration [41]. (b) Experimental schematic. A single sender strain containing an address-editable message plasmid is coupled with one of two receiver strains in pairwise transfer experiments. Prior to editing, the message is addressed only to the B receiver, but after editing, the message is addressed only to the A receiver. (c) Measured transfer rate from the experiment described in (b). Dots show the values from three biological replicates measured on different days, and bars depict the geometric mean of these values.

An important property of this address editing system is that it can be executed unilaterally by the sender cell, such that *a message’s recipient list can be updated without any coordination with the receivers themselves*. This feature is once again only possible because our framework encodes a message’s recipient list into the message itself, rather than relying on channel-specific interactions between the message vector and the recipient cell.

To assess the efficacy of our address editing system, we constructed a single sender strain that contains a nonmobilizable plasmid encoding a salicylate-inducible TP901 integrase expression cassette and the B gRNA binding site flanked by TP901 attB sites, alongside a message plasmid containing the A binding site flanked by TP901 attP sites in its address region. We then performed performed pairwise sender-receiver mating experiments with either of the two A- or B-expressing receiver strains from the original pairwise addressing experiments (Fig 3) in the presence or absence of salicylate induction (Fig 5b). As expected, transfer of the message to the A receiver was blocked in the absence of integrase activity (138-fold difference) while the blocking profile was reversed when the integrase was induced, with the transfer to the B receiver now being blocked (75-fold difference) (Fig 5c).

Comparing the transfer rates from the above experiment with the original pairwise transfer blocking experiments from Fig 3 reveals that the these latter data, which were measured in the absence of the address editing system, are nearly directly log-linearly related with the values obtained in the address editing experiment in all conditions except for the post-edit valid transfer (Fig S3). This transfer rate, which is supposed to be high, is 5-fold lower than expected from this correlation. It is unclear why this is the case, as factors like leaky integrase expression or an inability for the integrase to fully edit the message plasmid population would both manifest as deviations from predicted transfer rates in one of the blocked transfer conditions, neither of which occur in our results. It is therefore prudent to note that the address editing process can reduce the transfer rate to the new recipient cells through an unknown mechanism, although in this case the consequence was only a 2-fold reduction in the system’s dynamic range.

### Address editing enables control of information flow through a population

Having demonstrated that integrase-mediated cassette exchange can successfully edit the recipient list of a message plasmid, we next designed a linear message relay system to demonstrate how address editing can control the flow of information through a population. In this system, the message plasmid propagates in a linear sequence along a defined order of strains in a population, without skipping ahead or backtracking (Fig 6a). This sequential order is enforced by ensuring the message plasmid is only addressed to the next strain in the sequence at any given time, which can be implemented by having each successive strain edit the message’s address accordingly. An important but subtle property of this system is that the entire signal relay is implemented using a single communication channel that modifies its message at each step. Implementing a similar signal relay with SMA channels would require n − 1 orthogonal channels for an n-strain population, while the DNA-based implementation requires only a single channel regardless of the complexity of the consortium composition (Fig S4).

**Figure 6.**
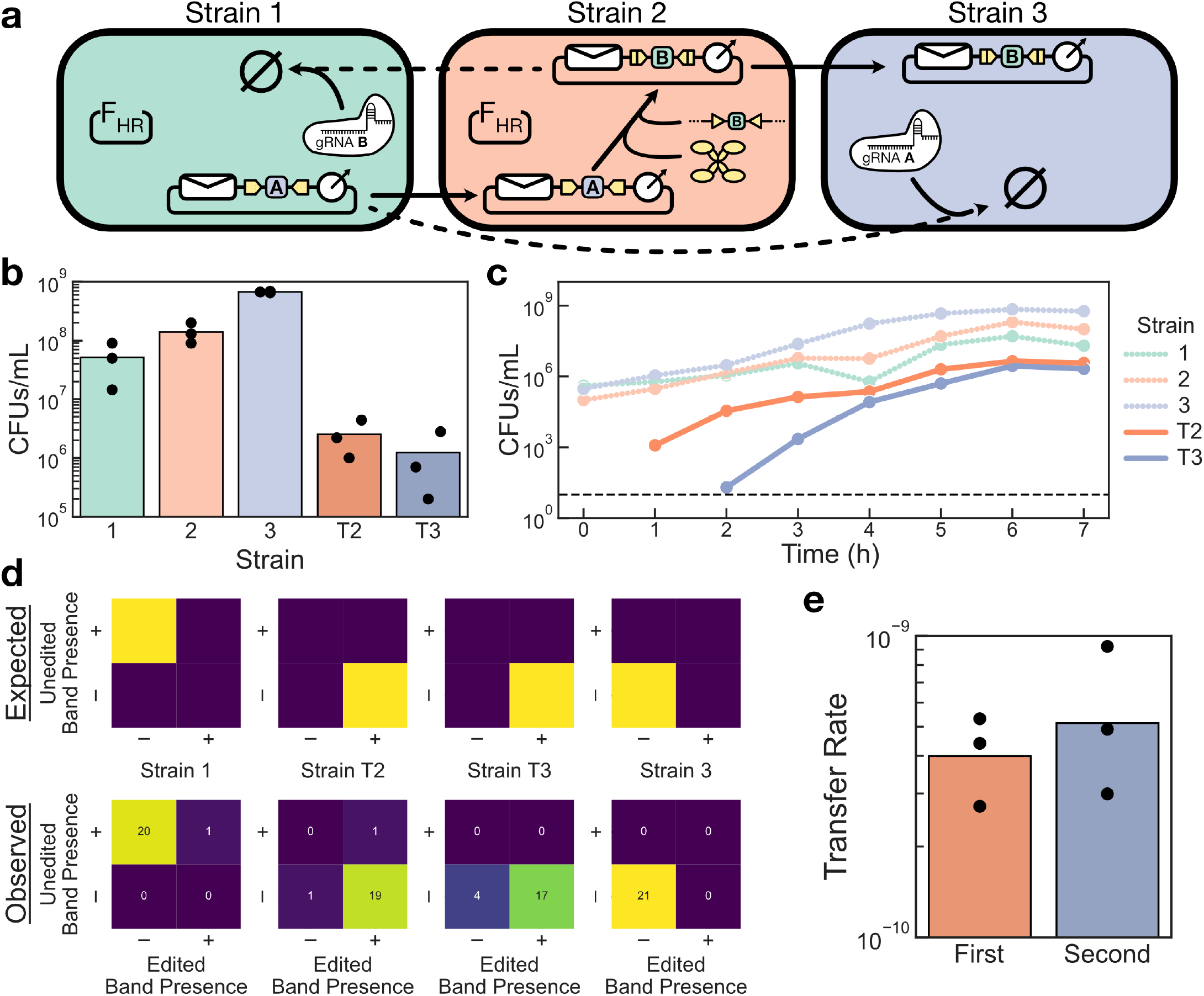
A three-strain linear message relay. (a) Schematic of the relay system. The message plasmid starts in Strain 1 and can only be transferred to Strain 2. When the message enters Strain 2, its address is edited so that it can no longer return to Strain 1 but is now allowed to continue on to Strain 3. This architecture is scalable to *n* strains (Fig S4) (b) Endpoint densities of each strain after 6 hours of coculture. (c) Timecourse plating showing the growth of each strain within the coculture over time for a single biological replicate. The dashed black line marks the limit of detection. (d) PCR assays of endpoint colonies from selected strains. (Top row) Based on the mechanism of the relay, different strains are predicted to contain either the edited or unedited version of the message. (Bottom row) Results of PCRs that selectively amplify either the edited or unedited version of the message from 21 colonies of each selected strain. The number of colonies that were assigned to each result condition are indicated in the heatmap. (e) Transfer rate calculated for each step in the relay, based on the data from (b). For the first transfer, Strain 1 is the sender and Strain 2 is the receiver. For the second transfer, Strain T2 is the sender and Strain 3 is the receiver. In all bar graphs, dots show the values from three biological replicates measured on different days, and bars depict the geometric mean of these values.

We designed the strains for a three-strain linear relay, as described in Fig 6a— Strain 1 contains F_HR_ and the message plasmid (initially addressed only to Strain 2) as well as the B gRNA, while Strain 2 contains F_HR_ and the machinery to edit the message plasmid, when received, to address only Strain 3. Strain 3 itself simply expresses the A gRNA. A unique antibiotic resistance gene was genomically-integrated into each strain (gentamicin, apramycin, and spectinomycin, respectively). We then mixed the three strains together and cocultured them for 6 hours before selectively plating out each strain (three parent strains and two possible transconjugant strains) to measure their densities in the endpoint population state (Fig 6b). Following expectations, the final density of Strain 3’s transconjugants (T3) was lower than the density of Strain 2’s transconjugants (T2), even though the total endpoint density of Strain 3 was 5-fold higher than that of Strain 2.

To further validate our system’s performance, we performed a timecourse assay for one of our replicates where we plated out the coculture every hour after the initial mixing to obtain the growth curves of each strain over the course of the experiment (Fig 6c). These results are again consistent with the desired system behavior, as Strain 3’s transconjugants do not appear until after Strain 2’s transconjugants have become detectable.

Finally, we performed PCRs on colonies from the endpoint cultures that selectively amplify either the edited or the unedited form of the message plasmid, which allows us to determine whether each strain contains only the expected form of the plasmid. The results aligned almost directly with the expectation for each tested strain, with the exception of four Strain 3 transconjugants showing sub-threshold amplification of both plasmid types (Fig 6d).

We additionally noted that the transfer rate of the second step in the relay was not lower than the transfer rate of the first step in the relay (Fig 6e), suggesting that editing a message plasmid’s address does not necessarily impose a detectable penalty on its transfer rate as was observed in Fig 5.

Taken together, these results confirm that the address editing system can indeed be used to reliably control the flow of messages through a population.

## Discussion

In this work, we have designed a modular, scalable, and adaptable message addressing framework for DNA-based communication channels and implemented it in an F-mediated plasmid conjugation system. Because our addressing system is built with molecular components that are orthogonal to the native horizontal gene transfer machinery, any existing DNA-based communication channel can be modified to incorporate our addressing system by expressing Cas9 and a strain-identifying gRNA in the receiver cells and encoding an address region onto the message vector.

Because our emphasis was to provide a proof-of-concept demonstration of the capabilities of a DNA communication system that fully takes advantage of the dynamic mutability of its messages, there are many fruitful directions for further optimization of this framework. For example, because we expressed our transfer blocking cassette from a plasmid in our receiver cells, it is likely that mutation and plasmid loss created a subpopulation of receivers without a functional transfer blocking cassette [43, 44]. By promoting its evolutionary stability, for example by integrating it onto multiple sites on the genome, it is possible that we could improve the system’s ability to block off-target transfers even further.

Another promising direction is to improve and augment the transfer properties of the original horizontal gene transfer system itself. F_HR_, like the M13 helper system, constitutively expresses its transfer machinery, but the master transcriptional regulators for these operons have been identified and so could be engineered to increase their expression or place them under inducible control [45]. Interfacing more with the system’s channel-intrinsic properties, for example by modulating the expression of entry exclusion proteins to globally block plasmid receipt [46], could also add an additional layer of programmable functionality to the system.

Converting additional horizontally-mobile genetic vectors into new DNA-based communication channels will also be an important component of the continued development of DNA messaging. For example, the F plasmid is known to stop conjugation as the population approaches stationary phase, which limits its overall transfer rate in liquid culture experiments [47]. We observed that this property can also hold for plasmids mobilized by F_HR_ (Fig S5) and that this leads to a low overall transfer rate— only around 50% of the receivers in our pairwise transfer experiments were converted to transconjugants after 6 hours of coculture in the absence of transfer blocking (Fig S6). In contrast, Ortiz and Endy were able to achieve over 90% receiver conversion after 5 hours of coculture using their M13 bacteriophage-based system [14], despite the fact that the M13 transfer rate has been estimated to be lower than the F transfer rate in coculture conditions [48]. As different applications will be best served by systems with different transfer properties, developing a diverse and well-characterized toolbox of DNA communication channels will be important in facilitating their wider use.

One potential class of applications where the use of our addressing system may not always be appropriate, however, is in cases where a transient amount of off-target expression would be detrimental. Because our system blocks transfer by degrading the message after it has entered the recipient cell, it is possible that genes on the message could be expressed in an off-target recipient before the message is cleaved and degraded— indeed, some genes carried on the F plasmid have been observed to express as soon as 10 minutes after the plasmid’s initial entrance into a receiver cell [49]. Preliminary experiments with our F_HR_ -based system, however, suggest that when genes are expressed weakly from the message plasmid, Cas9-mediated cleavage can occur quickly enough to prevent any detectable activity of these genes within off-target recipients (Fig S7). A thorough analysis of this phenomenon will likely require a comprehensive characterization of various transfer and blocking systems.

This transient expression phenomenon highlights the fact that DNA-based communication will not necessarily be the appropriate tool for every application. Indeed, although SMA channels do not exhibit many of the useful advantages of DNA channels, their simplicity and reliability nonetheless lets them fill a valuable niche for the efficient implementation of low-complexity communication. In contrast, the role that DNA messaging is well suited to play in the continued development of consortium engineering is to push the boundaries of achievable complexity in the space of behaviors that can be programmed into a system.

By leveraging the dynamic mutability of DNA messages, alongside the message-channel decoupling and high information density already demonstrated by Ortiz and Endy, our work serves as a “second step” in the foundation of DNA messaging by creating a single generalizable system that embodies all three of its unique advantages. The ability to leverage a decade of intensive efforts to develop effective molecular DNA editors was critical in enabling our framework, and as these tools continue to advance, DNA messaging is itself poised to increase its functional capacity. Such future progress in DNA messaging that improves and expands upon the three advantages highlighted in our system, for example by developing ways for cells to generate biologically-interpretable message *de novo*, will bring the field increasingly closer to realizing the ability to engineer autonomous, self-adapting multicellular systems that rival the complexity of living systems.

## Methods

### Strain and plasmid construction

A list of all strains and constructs used in this study, associated with the experiments in which they were used, can be found in the Supplementary Materials. The parent strain of the Keio single-gene knockout collection [50], *E. coli* BW25113, was used as the basis for all experiments in this study with the exception of those described in Supplemental Figures S5 (*E. coli* JS006 [51]) and S7 (*E. coli* Marionette MG1655 [52] for the receiver strains). Genomic integrations were performed using the pOSIP clonetegration system [53].

Because the F_HR_ plasmid retains a low rate of self-transfer activity and carries a tetracycline resistance gene, the plasmid could be transferred from the original F_HR_ donor strain into newly-constructed sender strains using standard mating procedures (see below) and selectively plating for transconjugants.

All new plasmids for this study were constructed via 3G assembly [54] using genetic parts from the CIDAR MoClo extension part kit [55, 56] when available. Parts not in the kit were converted to 3G-compatible parts by amplifying them with custom primers or synthesizing the parts directly before combining them with the part plasmid backbone via Gibson assembly. The former approach was used for the inducible promoters and their cognate regulators, taken from the Marionette system [52]; the spectinomycin and apramycin resistance genes, taken from the pQCascade and pCutAmp plasmids, respectively [57]; thee gentamycini resistance cassette, taken from the pJM220 plasmid [58]; and the F *oriT* sequence, taken from the mobile GFP plasmid developed by Dimitriu et al. [16].

The latter synthesis-based approach was used for the address regions and the gRNAs. The sequences for the “A” and “B” gRNAs were designed from the second half of the UNS2 sequence [59] and the recognition site of the I-SceI endonuclease [60], respectively.

All message plasmids in this study, with the exception of those in Fig S7, were constructed on a pSC101-origin backbone. The message plasmids in Fig S7 were constructed on a ColE1-origin backbone. All nonmobile plasmids used in this study were constructed on a p15a-origin backbone.

### Cell culturing and plasmid transfer experiments

Strains involved in the transfer experiments were grown overnight in 2mL of LB media in a 15mL polyproylene culture tube in a shaking incubator set to 37C and 250rpm under antibiotic selection for each resistance present in the strain. In the morning, each culture was diluted 1:100 into 2 mL of fresh LB media containing antibiotic selection for only the plasmid-based resistances in the strain and returned to the shaking incubator until the culture reached midlog phase (approx. 1-2 hours). At this point, cultures were induced with the appropriate amount of inducer, if applicable, and continued incubating for another 1 hour.

Cultures were then removed from the incubator and their OD600 value was measured. 1 mL of the culture was then transferred into a 1.5 mL tube and spun at 4,000 rpm for 10 min on a tabletop centrifuge. The supernatant was removed and the cell pellet was resuspended in 1 mL of fresh LB containing only kanamycin, alongside the appropriate concentration of inducer if applicable. Cultures were then spun again at the same settings and resuspended in fresh media as before.

Cells in the washed cultures were then diluted into a single 3 mL culture of fresh LB, containing only kanamycin and appropriate inducers, in a 15 mL culture tube. Cells were added such that each strain would have an OD600 value of 0.002 within the final coculture, which typically involved at least a 1:100 dilution from the original monoculture. The coculture was then placed back into the shaking incubator, marking the beginning of the 6 hour coculturing window.

The concentrations of the antibiotics in the medias, when used, were 25 μg/mL kanamycin, 12.5 μg/mL chloramphenicol, 25 μg/mL apramycin, 25 μg/mL spectinomycin, 15 μg/mL gentamicin, 50 μg/mL carbenicillin, and 10 μg/mL tetracycline.

### Selective plating and measuring strain densities

After 6 hours of coculturing (or at hourly timepoints, for Fig 6c and S5), the culture tube was removed from the incubator and the density of each strain was assessed by selective plating of serial dilutions of the culture. 60mm LB agar plates containing the appropriate antibiotic markers (the unique genomically-integrated resistance cassette for the strain as well as chloramphenicol to select for the message plasmid, when appropriate) were used for selection. For all experiments except those in Figs 6 and S5, serial dilutions were performed with 10-fold steps in 100 μL volumes and four 5 μL spots of successive dilutions spanning the expected density were spread onto a single plate. For the remaining experiments, serial dilutions were performed with 100-fold steps in 1 mL volumes and 100 μL of one or two dilutions was spread onto individual plates. Plates were incubated at 37C until the formation of colonies (12-24 hours).

Strain densities, measured as colony forming units per mL, were calculated by counting the number of colonies on the selection plates and multiplying by the appropriate dilution factor. When multiple dilution factors displayed growth, the dilution factor with the highest number of colonies that still remained countable (i.e. colonies were clearly discernible and separable) was used to calculate the density. Colonies were counted manually.

When fluorescent proteins were used to distinguish different colonies on the same selection plate, as in Fig 4, plates were imaged on an Olympus MVX10 microscope with the appropriate fluorescence filters.

### Calculating transfer rates

The transfer rate was calculated as T/(S* R), where T is the density of transconjugants, S is the density of senders, and R is the *total* density of receivers. Note that unlike some recent works that use this transfer rate measure [61, 62], we choose to define the value of R to include the density of transconjugants, rather than representing the density of receivers without the transferred plasmid. This choice was made so that experiments where transconjugants could not be discerned from receivers on receiver-selecting plates, such as those in Fig 4, could be analyzed in the same way as those where the distinction could be made, and so that if the transfer ever went to completion and all receivers were converted into transconjugants, the value of the transfer rate would not be undefined. The fractional receiver conversion value, as shown in Fig S6, can then be calculated as T/R.

### PCR assay for message plasmid identity

Colonies were picked and resuspended into 10 μL of M9 minimal media, of which 1 μL was placed into two separate 10 μL PCR reactions with primers designed to bind to the *oriT* and either the A or B gRNA binding site. Primer sequences were, for the unedited message plasmid (A site), CGCAGAATCCAAGCCG and CGGATAAAGTCACCAGAGGTG (with an annealing temperature of 64C) and for the edited message plasmid (B site), GGGATAACAGGGTAATC and GATAAAGTCACCAGAGG (with an annealing temperature of 56C).

The number of PCR cycles was adjusted for each reaction against positive control colonies (cells containing a single message plasmid with either just the A site or just the B site on its address) to reduce the probability of observing false positives in the assay. The temperature program was 5 minutes at 98C followed by N cycles of 10 seconds at 98C, 30 seconds at the annealing temperature, and 20 seconds at 72C. After the cycles were completed, the reaction was kept at 72C for an additional 5 minutes before cooling down to 4C. N was 23 for the reaction targeting the unedited address and N was 26 for the reaction targeting the edited address.

PCR samples were run on a gel and imaged on a UV imager, and the presence and absence of a band for each sample was determined by eye.

## Supporting information

CFU count data

Strain and Construct List

## Data and Strain Availability

Raw data for CFU counts can be found in the Supplementary Materials. As of this writing, strains and plasmids created for this study are being prepared for deposition to AddGene. Sequence maps for the constructs created for this study are available upon request.

## Acknowledgments

We thank A. Halleran, A. Shur, and R. Williams for insightful discussions and the members of the Murray lab for feedback on the manuscript. We also thank T. Dimitriu for providing plasmids from the F_HR_ system. This material is based upon work supported by the National Science Foundation Graduate Research Fellowship Program under Grant No. DGE-1745301 and by the Institute for Collaborative Biotechnologies through grants W911NF-09-D-0001 and W911NF-19-2-0026 from the U.S. Army Research Office. The content of the information in this report does not necessarily reflect the position or the policy of the Government, and no official endorsement should be inferred.

## Author Contributions

J.P.M. conceived and designed the project, performed the experiments, analyzed the data, and wrote the manuscript, all under the supervision of R.M.M.

## Supplementary Figures

**Figure S1.**
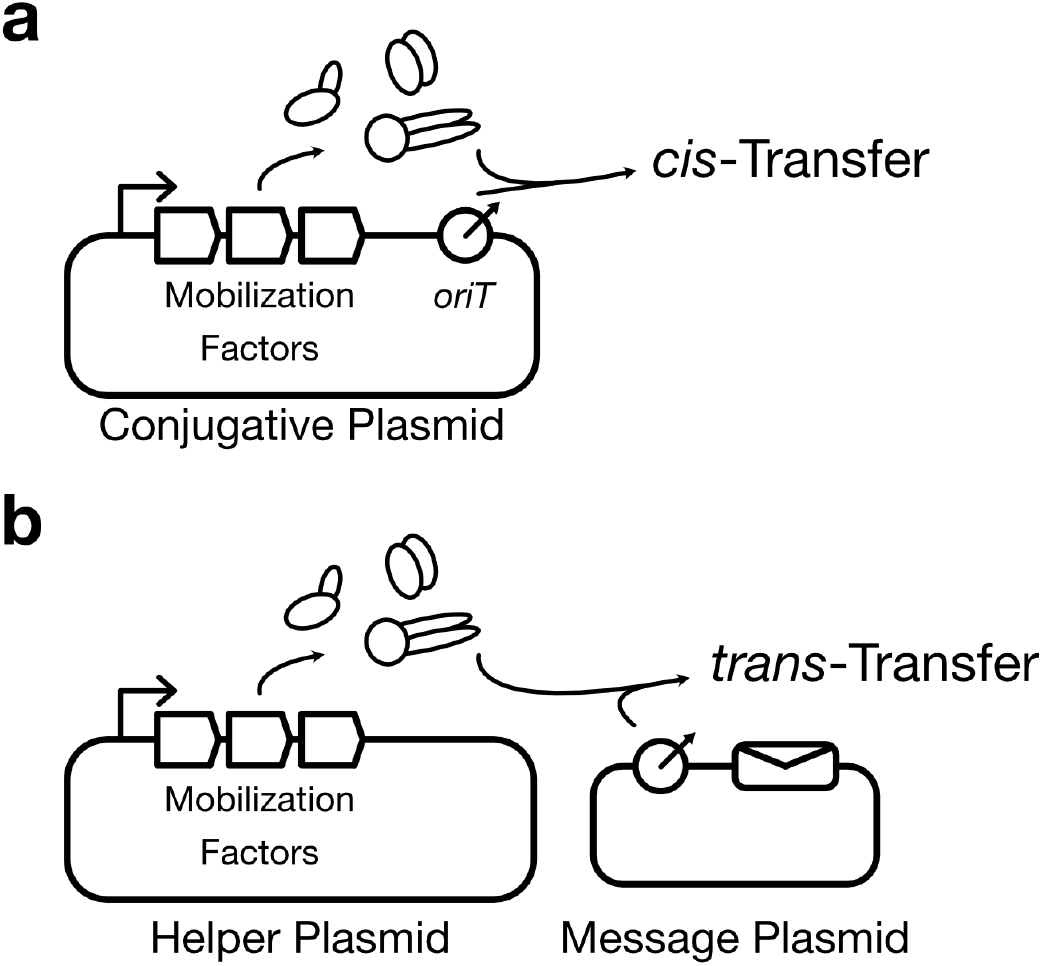
How natural horizontal gene transfer systems are converted into DNA messaging channels. (a) Schematic of the architecture of a natural horizontal gene transfer system, using a conjugative plasmid as an example. The mobile vector expresses a set of genes, collectively called the mobilization factors, that transfer DNA elements that contain a cognate recognition sequence called the origin of transfer (*oriT*). Because the conjugative plasmid itself contains an *oriT* site, it transfers itself in a process termed *cis*-trasnfer. (b) Schematic of the architecture of a DNA messaging channel. The *oriT* is removed from the conjugative plasmid to create a helper plasmid that confers the ability to transfer DNA to its host cell but cannot transfer itself. The cognate *oriT* sequence can be placed onto another DNA vector to create a DNA message, which can then be transferred to another cell via the mobilization factors expressed by the helper plasmid. This process is called *trans*-transfer. Other horizontal gene transfer mechanisms, like non-lytic bacteriophages, share this same fundamental architecture and can be converted into DNA messaging channels through this same process.

**Figure S2.**
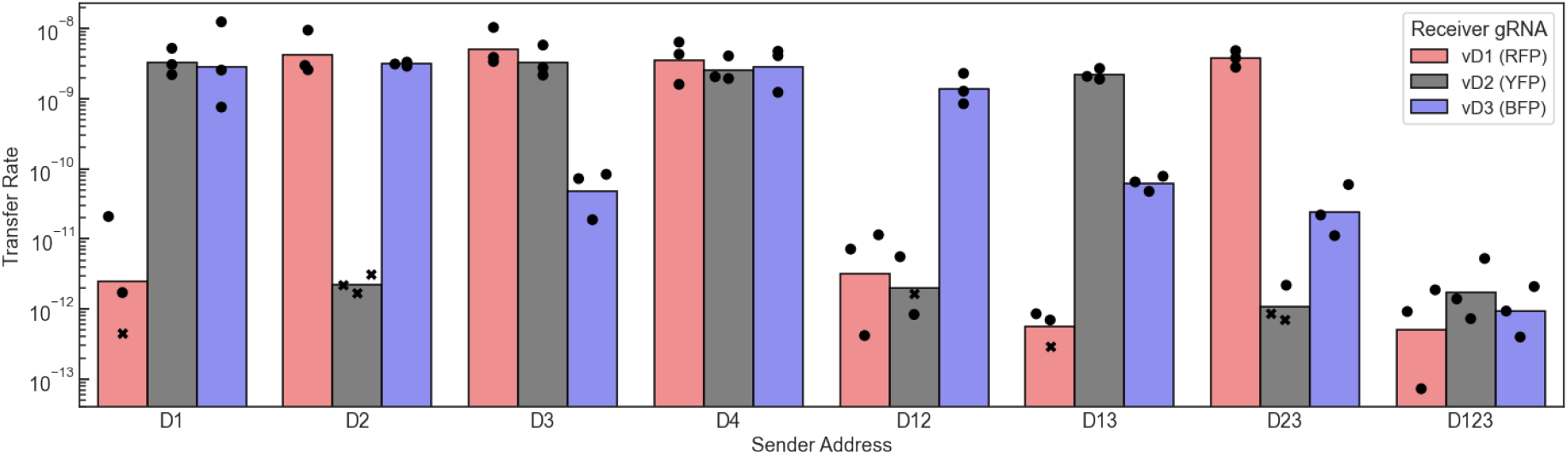
Transfer rate values for each biological replicate from Fig 4. In some conditions, the transconjugant densities were below the limit of detection (200 CFU/mL). In such cases the subthreshold strain density was assumed to be equal to the limit of detection for the purposes of calculating the transfer rate. Rates where this occurred are marked with an X instead of a circle, and are overestimates of the true transfer rate. Bars represent the geometric mean of the three biological replicates.

**Figure S3.**
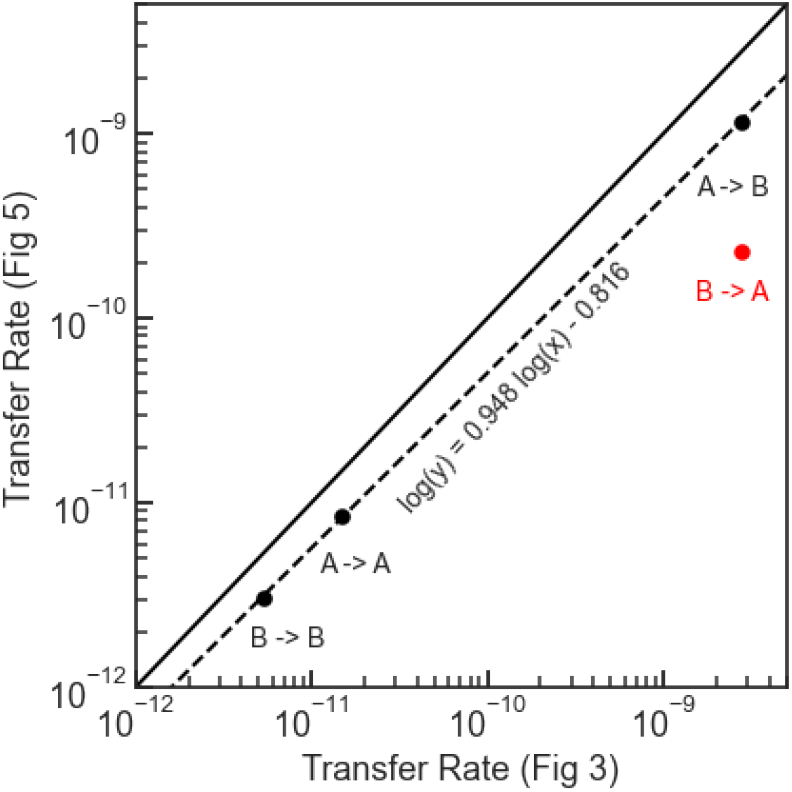
Geometric mean transfer rate values for the pairwise transfer blocking experiments from Fig 3 and Fig 5. Each transfer is marked as “I -> J”, where I is the site on the address region and J is the gRNA in the receiver. Integrase-induced conditions from Fig 5 are written using the post-edit site on the address region. The solid line shows a direct log-linear relationship log(y) = log(χ), while the dotted line is the result of a log-linear fit to the three black points. The vertical distance between the red point and the dotted line is 5.2-fold.

**Figure S4.**
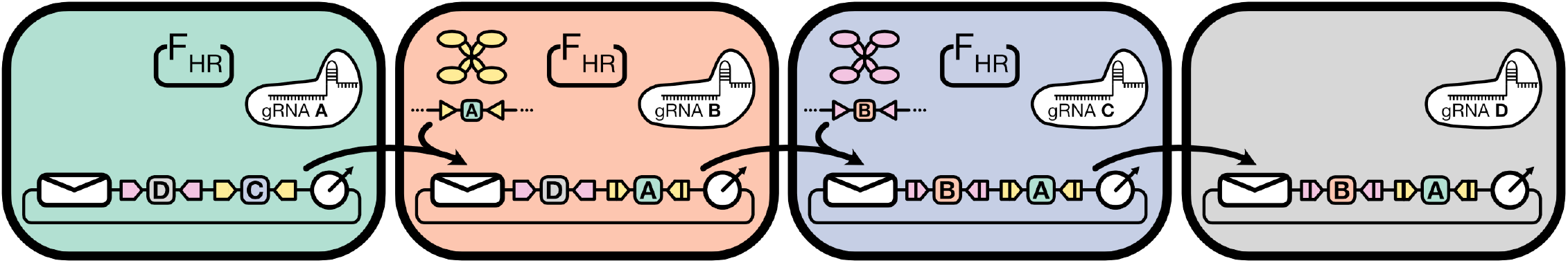
Schematic of a four-strain linear relay, which has the same architecture as an n-strain relay for n ⩾ 4. Each strain in the population expresses one of n orthogonal gRNAs, and the address region on the message plasmid contains n − 2 binding sites that block its transfer to all strains except its current strain and the next strain in the relay. Each site on the address is flanked by one of n − 2 orthogonal integrase attachment site pairs. All strains except the last strain in the relay contain F_HR_, and all strains except the first and last strains express a unique integrase that performs an address editing operation that invalidates the previous strain while validating the next strain in the sequence. Blocked transfers and intermediate message plasmid states are omitted from the diagram.

**Figure S5.**
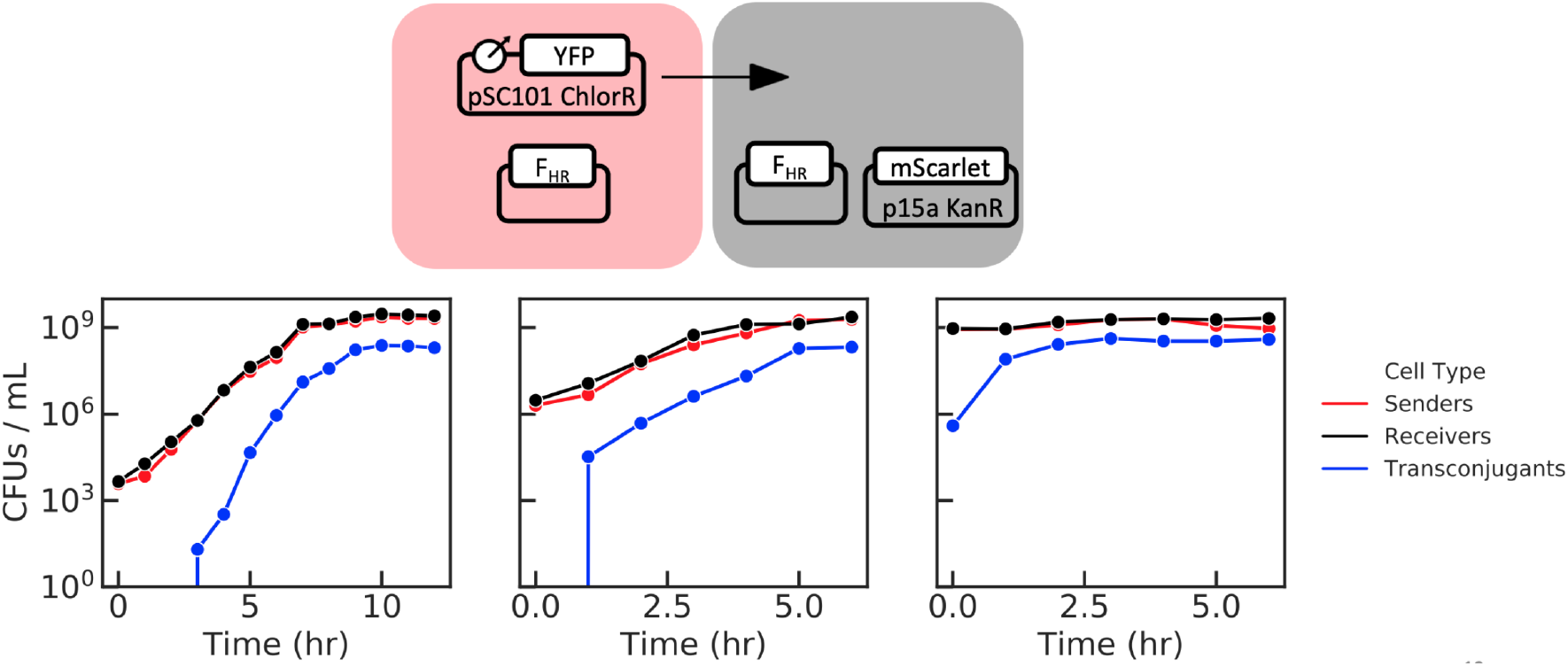
Timecourse plating results of F_HR_ -mediated mating experiments without the transfer blocking system, performed in *E. coli* JS006 cells, conducted in shaking LB media (as described in the Methods) without antibiotics. Selective plating with chloramphenicol alone, kanamycin alone, or both antibiotics together was used to calculate the total sender, receiver, and transconjugant density, respectively. The three graphs represent three distinct biological replicates, each one having a different initial strain density for the coculture. In each case, the transconjugant density plateaus before overtaking the entire population.

**Figure S6.**
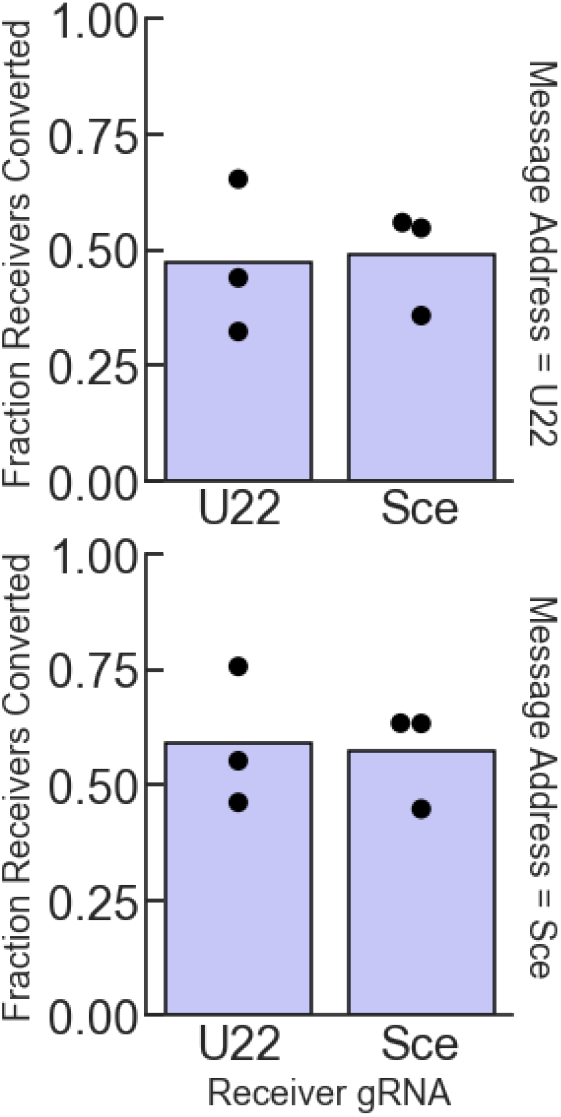
Fraction of receivers converted to transconjugants, T/R, in the Cas9-uninduced conditions of the transfer experiments from Fig 3. The mean value across all conditions is 53%.

**Figure S7.**
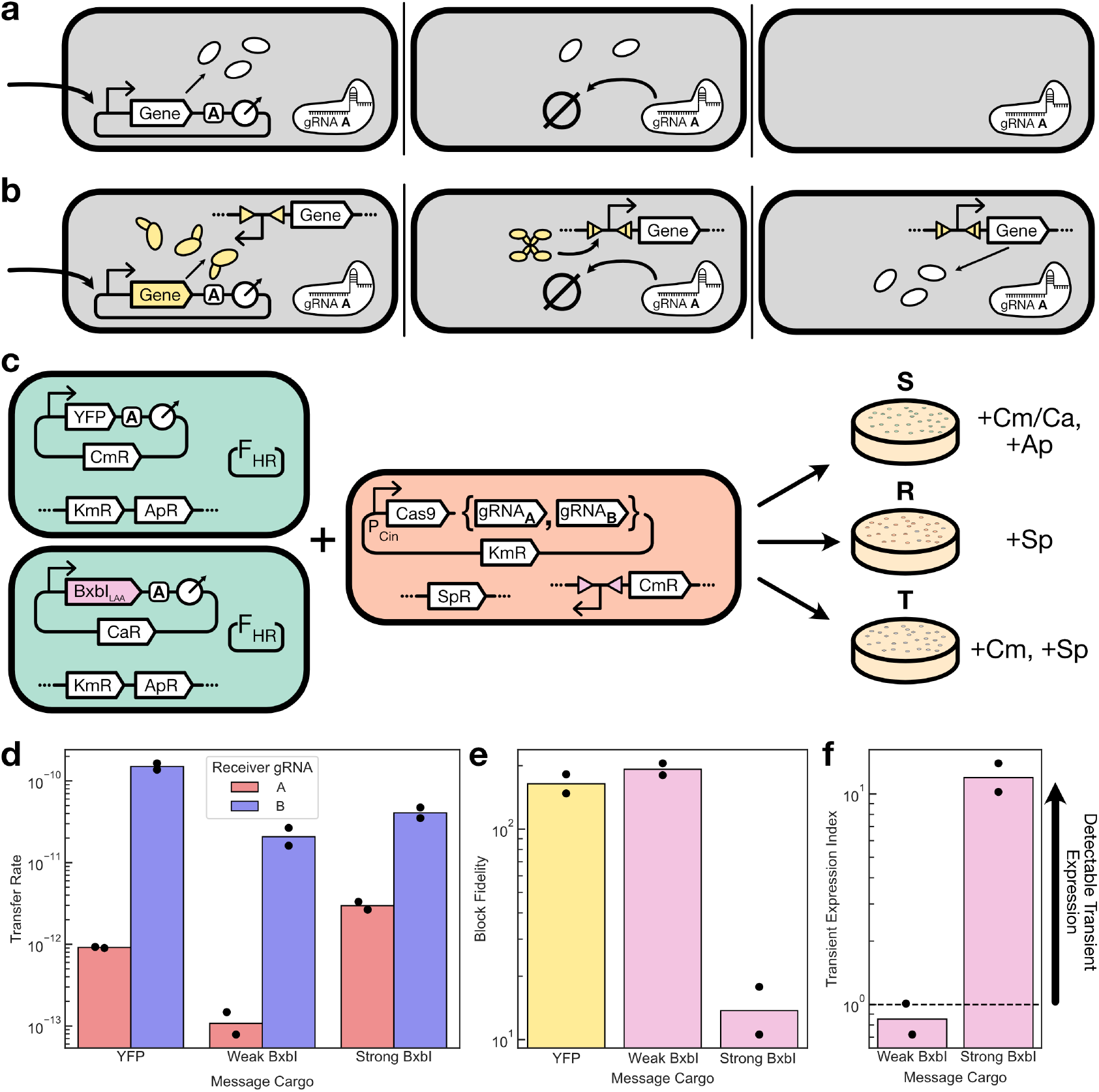
(a) Schematic of transient expression from a blocked message. (Left panel) An invalid message enters the recipient cell and begins expressing genes. (Middle panel) The Cas9-gRNA complex degrades the message plasmid, but the expressed proteins remain in the cell. (Right panel) Eventually, these proteins are degraded or diluted out of the cell and the influence of the invalid message disappears. (b) Schematic of transient expression inducing a permanent change in the receiver cell. (Left panel) An invalid message plasmid expressing an integrase enters a cell that contains the cognate attachment sites. (Middle panel) The integrase can quickly act on the attachment sites and induce a permanent genetic change in the receiver cell. (Right panel) Even long after the invalid message has left the cell, its influence persists. (c) Experimental schematic. Two types of message plasmids were constructed: one carrying a constitutive YFP and chloramphenicol resistance cargo, and one carrying a degradation-tagged BxbI integrase and carbenicillin resistance cargo. These were mixed in pairwise transfer experiments with MG1655 Marionette-based receiver strains [52] expressing a cognate or non-cognate gRNA and a BxbI-activatable chloramphenicol resistance cassette integrated onto the genome. Transconjugants were always selected with chloramphenicol, so that the YFP plasmid transfer rate captures only transconjugants that currently contain the message plasmid while the BxbI plasmid transfer rate captures receiver cells that at one point in their ancestry received the plasmid for sufficient time to express BxbI and activate the chloramphenicol resistance cassette. (d) Measured transfer rates for the YFP and BxbI plasmids. Two variants of the BxbI plasmids, identical except for their expression strength, were tested. (e) The block fidelity, defined as the ratio of the transfer rates between the valid and invalid receiver, for the experiments. Higher values indicate better blocking. (f) The transient expression index, defined as the ratio of block fidelity between the BxbI plasmid and the YFP plasmid. A value > 1 indicates that the YFP plasmid was blocked more effectively than the BxbI plasmid, suggesting that transient expression of the BxbI in invalid receiver cells inflated the endpoint transconjugant density detectably. A value ⩽ 1 indicates that transient expression of BxbI in the invalid receiver cells was not detectable in this assay. Dots represent two biological replicates measured on different days, and bars indicate their geometric mean.

